# Human Dermal Fibroblast-derived Spheroids Demonstrate Efficacious Immune Modulation in a Psoriasis Mouse Model

**DOI:** 10.1101/2025.08.04.668478

**Authors:** Chuo Fang, Inah Bianca Embile, Christen Boyer, Simon Gebremeskel, Nikolay Bazhanov, Bin Jiang, Subhiksha Raghuram, Midori Taruishi, Pete O’Heeron, Hamid Khoja

## Abstract

**Background:** Psoriasis is a chronic inflammatory disease associated with high morbidity and few cases of sustained remission. Innovative immunomodulatory therapies, including fibroblast-based cell therapies, offer promising alternatives. This study investigates the therapeutic potential of human dermal fibroblasts (HDFs) organized into three-dimensional (3D) spheroids in a mouse model of imiquimod (IMQ)-induced psoriasis.

**Methods:** HDF spheroids were cultured using Elplasia® microcavity plates, and their size, viability, and phenotype were compared with single cells in 2D monolayer cultures. Cellular responses in whole blood and acute inflammatory responses were evaluated at various time points following intravenous injection of HDFs. The therapeutic efficacy of HDF spheroids was assessed using an IMQ-induced psoriasis mouse model, with disease severity scored using the Psoriasis Area and Severity Index (PASI). Optimized HDF spheroids (∼150 µm, 1×10^6^ cells/mouse) were administered intravenously in a single dose for mild psoriasis or multiple doses for moderate-to-severe psoriasis. The efficacy of HDF spheroids was compared to a pre-clinical monoclonal antibody targeting interleukin 23 (anti-IL-23).

**Results:** Spheroid cultures of HDFs showed reduced cell size, enhanced viability, and distinct phenotypic changes compared to monolayer cultures. Intravenous injection of HDF spheroids resulted in less thrombocytopenia and reduced acute inflammatory responses compared to single-cell injection. A single dose of HDF spheroids reduced the severity of mild psoriasis by 35%, while repeated doses resulted in a 36% reduction in moderate-to-severe psoriasis. Single-dose administration normalized blood cell counts, alleviated spleen enlargement, and improved cytokine dysregulation. Although HDF spheroids and anti-IL-23 reduced epidermal thickening and immune cell infiltration, HDF spheroids uniquely inhibited monocyte production and infiltration, a benefit not observed with anti-IL-23. No acute or chronic toxicity was observed.

**Conclusions:** HDF spheroids offer comparable therapeutic efficacy to anti-IL-23 in treating psoriasis, with a distinct mechanism involving inhibiting monocyte production and infiltration. Their safety profile and broader immunomodulatory potential support their development as a novel therapeutic strategy for psoriasis and other inflammatory diseases.

## Background

Psoriasis is a chronic skin inflammatory condition affecting approximately 2-3% of the global population across all age groups. It is characterized by hyperproliferation of keratinocytes, leading to the formation of red, scaly patches on the skin that significantly impact patients’ quality of life. The severity of the disease can range from mild to severe, with the involvement of extensive areas such as the scalp, elbows, knees, and nails (1). Additionally, psoriasis is associated with considerable comorbidities, including psoriatic arthritis, cardiovascular diseases, and metabolic syndrome, further complicating its impact on public health (2). Despite advances in treatment, current therapies often provide inadequate long-term relief and may result in substantial adverse side effects, highlighting the need for novel and more effective therapeutic options (3).

The pathophysiology of psoriasis is complex, involving sustained inflammation that leads to uncontrolled keratinocyte proliferation and dysfunctional differentiation. This inflammation arises from disruptions in both innate and adaptive immune responses, affecting not only the epidermal layer but also the interactions between keratinocytes and various cell types throughout the dermal layer. Activated T cells, particularly Th1 and Th17 subsets, release pro-inflammatory cytokines such as tumor necrosis factor α (TNF-α), interleukin (IL)-17, and IL-22, which exacerbate inflammation and contribute to the formation of psoriatic plaques (4, 5). Although targeted therapies, including biologics that block TNF-α, IL-17, and IL-12/23 signaling, have shown significant efficacy, they are accompanied by adverse side effects, variable responses, diminishing effectiveness over time, and the need for long-term treatment (6, 7). Cell therapy is emerging as a promising alternative, with mesenchymal stem cells (MSCs) demonstrating potential in alleviating psoriasis severity (8–12). MSCs are believed to exert their effects through their immunomodulatory and anti-inflammatory properties, influencing various immune cells and directly influencing keratinocytes, T lymphocytes, macrophages, and dendritic cells (DCs) to mitigate disease symptoms (8–10, 12).

Fibroblasts are increasingly recognized as a promising cell therapy for autoimmune diseases due to their phenotypic and functional similarities with MSCs and their notable immunomodulatory properties (13). In clinical settings, fibroblasts have demonstrated effectiveness in wound care, including the treatment of diabetic foot ulcers (14) and recessive dystrophic epidermolysis bullosa (15). Their therapeutic potential extends to pre-clinical models of autoimmune diseases, such as type I diabetes (16), alopecia areata (17), arthritis (18), and multiple sclerosis (19). These studies highlight fibroblasts’ ability to modulate the immune response by stimulating regulatory T cells (Tregs), suppressing pro-inflammatory Th17 cells, and inhibiting DC maturation (16–19). Furthermore, fibroblasts address the scalability and cost challenges often associated with MSC therapies (13), making them a potentially cost-effective alternative, particularly for patients requiring long-term treatment or at risk of relapse. Despite these promising attributes, the therapeutic potential of fibroblasts in experimental psoriasis models has yet to be explored.

Emerging evidence suggests that aggregating MSCs into three-dimensional (3D) spheroids significantly enhances their therapeutic efficacy in treating various inflammatory diseases. This approach has shown promise in murine models of wound healing (20), peritonitis (21), inflammatory bowel disease (22), inflammatory pulmonary disease (23), and peripheral artery disease (24), as well as in non-human primate models of multiple sclerosis (25) and osteoarthritis (26). Based on these findings, we hypothesized that 3D spheroids formed from human dermal fibroblasts (HDFs) could serve as an effective cell-based therapy for psoriasis. To test this hypothesis, we comprehensively characterized HDF spheroids, assessing their size, viability, and phenotypic features in 3D culture. Additionally, we explored their therapeutic potential in a clinically relevant mouse model of psoriasis induced by topical imiquimod (IMQ), which activates immune cells and triggers the release of pro-inflammatory cytokines, particularly those involved in the IL-23/IL-17 axis (27). The efficacy of HDF spheroids was compared with a monoclonal antibody targeting IL-23 (anti-IL-23), revealing that HDF spheroids offer a distinct mechanism by mitigating psoriasis-related inflammation through the suppression of macrophage production and infiltration in affected tissues, a benefit not observed with anti-IL-23.

## Methods

### HDF spheroid culture

HDFs purchased from Gibco (Thermo Fisher Scientific, Waltham, MA, USA) were cultured in Cellartis MSC Xeno-Free Culture Medium (Takara Bio, Japan) at 37°C with 5% CO_2_ until they reached 70-80% confluence. For 3D spheroid culture, Elplasia® ultra-low attachment microcavity plates and flasks (Corning, New York, USA) were used according to the manufacturer’s instructions. HDFs (passages 5-7) were seeded into 24-well Elplasia® plates, with 125, 250, 500, 1,000, and 2,000 cells in each spheroid. After 4 days, spheroids were harvested and washed twice with phosphate-buffered saline (PBS) prior to further experiments. Spheroids were imaged at 5 representative areas (totaling 45-54 spheroids) with an Echo Revolve Microscope (Echo, San Diego, California, USA) and characterized by measuring their diameter (μm) and the cross-sectional area (μm^2^) using ImageJ software version 1.53 (NIH, Bethesda, MD, USA). To assess phenotypic differences between HDF single cells and spheroids, cells were incubated with 0.25% trypsin-EDTA (Thermo Fisher Scientific, Waltham, MA, USA) for 5-15 min depending on the size at 37°C. The dissociated cells were re-plated (5×10^4^ cells/well) on 2-well Nunc Lab-Tek II CC2 chamber slides (Thermo Fisher Scientific, Waltham, MA, USA) and prepared for immunofluorescent staining.

### Cell viability in HDF single cells and spheroids

To evaluate if single cells aggregating into spheroids would impact cell viability, HDF single cells and spheroids were suspended in PBS supplemented with 10% fetal bovine serum (FBS) and stored at 4°C. After 7 days, HDF spheroids were collected and allowed to attach to a T-25 flask overnight. Afterward, spheroids were trypsinized at 37°C for 15 min with gentle agitation to facilitate the enzymatic breakdown of the extracellular matrix (ECM). During incubation, the spheroids were gently pipetted up and down to aid the dissociation process. The dissociated cells were assessed for viability and diameter using Via1-Cassettes and Solution 10 with the NucleoCounter NC-3000 instrument (Chemometec, Allerod, Denmark) according to the manufacturer’s instructions. Cell viability and diameter were compared between HDF single cells and spheroids.

### Pre-clinical safety evaluation of intravenous injection of HDF spheroids

In cell-based therapy, the interaction between transplanted cells and blood components (e.g., platelets) can trigger the instant blood-mediated inflammatory reaction (IBMIR), leading to serious complications, including microvascular thrombosis and ischemia (28). In order to assess the risk of IBMIR, C57BL/6J female mice (10 weeks of age, 20-25 g) purchased from The Jackson Laboratory (Bar Harbor, ME, USA) were randomly assigned (n=5/group) to receive Plasma-Lyte A (PLA, Baxter, Deerfield, IL, USA) as the vehicle control, HDF single cells (1×10^6^ cells/mouse), or HDF spheroids (1×10^6^ cells/mouse, 1,733 cells in each spheroid with an average diameter of 150 μm). Immune cells and coagulation-related changes and immune cell changes were monitored at 1 hour, 24 hours, 3 days, and 7 days post-intravenous injection.

To further assess the short-term and long-term safety of intravenous injection of HDF spheroids, mice administered with HDF spheroids (1×10^6^ cells/mouse, 1,733 cells in each spheroid with an average diameter of 150 μm) or PLA were assessed in two periods: an acute toxicity evaluation at 2 weeks post-injection and a chronic toxicity evaluation over 12 weeks. Endpoints included measurement of relative tissue weights (heart, lungs, kidneys, liver, spleen, and brain), blood analysis for complete blood count (CBC), and chemistry panel for liver and kidney functions (Baylor College of Medicine Pathology Core and Lab, Houston, TX, USA), and histological examination of lung tissues using H&E staining and Masson’s trichrome staining.

### Mouse model of psoriasis and treatment

C57BL/6J female mice (10 weeks of age, 20-25 g) were randomly assigned to receive IMQ with PLA or HDF spheroids via the tail vein. In order to induce psoriasis, mice were anesthetized with isoflurane, and a rectangular area of 3 cm x 4 cm on the back skin was shaved and treated with Nair™ hair-removal cream on Day 0. Psoriasis was induced via topical application of 31.25 mg of IMQ cream (5% IMQ, Taro Pharmaceutical, Hawthorne, NY, USA) to the lower dorsal skin for 7 **(Figure 3A)** or 14 consecutive days **(Figure 4A)** (29). Mouse body weights were recorded daily. For cell-based treatment, HDF spheroids cultured in 6-well Elplasia® plates (seeded at 5×10^6^ cells/well) or 12K flasks (seeded at 20×10^6^ cells/flask) were washed twice with PBS, resuspended in PLA, and 577 spheroids were administered via the tail vein at a dosage of 1×10^6^ cells/100 μL/mouse. All experimental procedures followed the NIH Guide for the Care and Use of Laboratory Animals and were approved by the Institutional Animal Care and Use Committee at the K2Bio Lab (Houston, TX, USA) and StillMeadow, Inc. (Sugar Land, TX, USA).

### Clinical assessment of psoriasis

The clinical features of IMQ-induced psoriasis were evaluated daily using the Psoriasis Area and Severity Index (PASI) as described previously (30). Briefly, double-folded skin thickness was measured using a digital micrometer (Rexbeti, Auburn, Washington, USA). Psoriatic skin lesions, including thickness, erythema, and scaling, were assessed by two observers daily on a scale of 0 to 4 (0-no incidence; 1-slight; 2-moderate; 3-marked; 4-severe), respectively. No significant deviation was noted between observers. The scores from two observers were then averaged to determine the mean PASI for each mouse. Single and multiple administrations (every 3 days) of HDF spheroids occurred at mild psoriasis (average PASI score 2) and moderate-to-severe psoriasis (average PASI score 3-4), respectively. In order to compare the therapeutic efficacy of HDF spheroids with a pre-clinical monoclonal antibody targeting mouse IL-23 (anti-IL-23, BioxCell, Lebanon, NH, USA), anti-IL-23 was administered intraperitoneally on Days 1, 3, and 5 (IMQ+Anti-IL-23) (31); A single dose of HDF spheroids (IMQ+HDF) was intravenously injected on Day 4 (average PASI score 2) as described previously, and euthanasia occurred on Day 8.

### Blood and tissue collection

Blood was collected from the left ventricle via cardiac puncture under anesthesia. Approximately 100 µL of blood was drawn into EDTA-coated microcentrifuge tubes (BD, Franklin Lakes, NJ, USA), and the CBC profile was determined at Baylor College of Medicine Pathology Core and Lab (Houston, TX, USA). An additional 250 µL of blood was collected into serum-separator tubes and allowed to clot at room temperature (RT) for 30 min. Approximately 10 mg of skin tissue was weighed and placed in 500 µL of lysis buffer containing RIPA buffer and a protease inhibitor cocktail (Thermo Fisher Scientific, Waltham, MA, USA). The tissue was homogenized thoroughly and incubated on ice for 30 min to allow complete lysis. The lysates were then centrifuged at 12,000 x g for 15 min at 4°C, and the supernatant was transferred to a new tube. Protein concentration was determined using the bicinchoninic acid (BCA) assay (Thermo Fisher Scientific, Waltham, MA, USA), following the manufacturer’s instructions. Serum IL-17F and skin IL-10 levels were determined with MILLIPLEX® Mouse Cytokines/Chemokines Magnetic Bead Panel using Luminex multiplex bead assay (Millipore Sigma, St Louis, MO, USA) according to the manufacturer’s instructions and data were analyzed using the xPONENT software.

For histological analysis, mouse skin and lung tissues were fixed overnight in 4% paraformaldehyde (PFA, Thermo Fisher Scientific, Waltham, MA, USA) and sequentially passed through 15% sucrose/1× phosphate buffer and 30% sucrose/1x phosphate buffer until tissue sank prior to embedding in OCT (Sakura, Torrance, CA, USA). Skin and lung sections (5 μm) were obtained using a cryostat (Leica Biosystems, Heerbrugg, Switzerland) and stained with H&E and Masson’s Trichrome according to the manufacturer’s instructions (Abcam, Waltham, MA, USA). To assess the epidermal thickness, 3 random brightfield H&E images were obtained for each section using Echo Revolve Microscope (Echo, San Diego, CA, USA) and analyzed using ImageJ software version 1.53 (NIH, Bethesda, MD, USA). The epidermal thickness was calculated by dividing the area of the epidermis by its length (32).

### Immunohistochemistry

HDF spheroids were fixed overnight in 4% PFA, sequentially passed through sucrose solution, and frozen in OCT as described previously. Spheroid sections (5 μm) were prepared using a cryostat, rehydrated, and blocked with 1% bovine serum albumin (BSA, Thermo Fisher Scientific, Waltham, MA, USA) in PBS for 1 hour at RT. Blocked sections were then incubated overnight at 4°C with primary antibodies for CD44, CD90, CD105, Actin, Vimentin, S100A4 (1:200, Abcam, Waltham, MA, USA), and integrin α5 (ITGA5) (1:200, Thermo Fisher Scientific, Waltham, MA, USA). Mouse skin sections (5 μm) were stained with primary antibodies (1:200) for CD3 (T lymphocytes, Abcam, Waltham, MA, USA), F4/80 (macrophages, Thermo Fisher Scientific, Waltham, MA, USA), Ki-67 (cell proliferation marker, Abcam, Waltham, MA, USA), and keratin 14 (K14 for basal keratinocytes, Biolegend, San Diego, CA, USA). Afterward, sections were incubated with secondary antibodies (1:1000, Thermo Fisher Scientific, Waltham, MA, USA) for 1 hour at RT. For CD3 and F4/80 staining, sections were incubated with avidin-biotin-peroxidase (ABC) complex (Vector Laboratories, Burlingame, CA, USA) for 45 min at RT, developed using 3,3′-diaminobenzidine (DAB) (Vector Laboratories, Burlingame, CA, USA), and counterstained with nuclear fast red (Vector Laboratories, Burlingame, CA, USA). For Ki67 and K14, sections were stained with DAPI (1:1000, Thermo Fisher Scientific, Waltham, MA, USA) for 5 min. Three random images were obtained for each section (Echo, San Diego, CA, USA). The immunopositive area (expressed as a percentage of the total analyzed area) was quantified using NIH ImageJ software 1.53 (29).

### Flow cytometry

Spleens were harvested and placed in cold RPMI-1640 medium supplemented with 10% FBS and 1% penicillin-streptomycin. Spleens were then digested in the medium containing 150 μg/mL Liberase (Millipore Sigma, St Louis, MO, USA) for 1 hour at 37°C and homogenized by gently pressing through a 70 µm cell strainer (Greiner Bio-One, Kremsmünster, Austria) using the plunger of a syringe. The homogenate was collected into a 15 mL conical tube, and the cell suspension was centrifuged at 300 x g for 7 minutes at 4°C. The supernatant was discarded, and the cell pellet was resuspended in 1 mL of RBC lysis buffer (Thermo Fisher Scientific, Waltham, MA, USA) and incubated for 3-5 minutes at RT. Cells were washed twice with PBS by centrifugation at 300 x g for 5 minutes at 4°C and resuspended in 1 mL of PBS containing 2% FBS. Cells were blocked with anti-mouse CD16/CD32 (TruStain FcX^TM^, BioLegend, San Diego, CA, USA) and stained with fluorophore-conjugated antibodies, including CD11b (BV421-conjugated, BD Biosciences, Franklin Lakes, New Jersey, USA) and F4/80 (Alexa Fluor 488-conjugated, Thermo Fisher Scientific, Waltham, MA, USA). Following 10 min incubation at 4°C in the dark, cells were washed twice with PBS and resuspended in 100 μL PBS containing 2% PFA. Stained cells were analyzed using a CytoFLEX flow cytometer (Beckman Coulter, Brea, CA, USA). The percentage of macrophages within the splenocyte population was determined based on F4/80 and CD11b, and data were analyzed using FlowJo software (Tree Star, Inc.). The absolute number of macrophages was calculated by multiplying the total number of splenocytes, determined by cell counting using a NucleoCounter NC-3000 instrument, by the percentage of macrophages identified through flow cytometry (33).

### Statistical analysis

Statistical analysis was performed using GraphPad Prism version 10 (GraphPad Software, La Jolla, CA, USA). Results are expressed as the Mean ± Standard Error of the Mean (SEM). Normality tests were performed, and statistical differences between groups were analyzed using one-way ANOVA followed by Holm-Sidak’s multiple comparison test for parametric data or Kruskal-Wallis test followed by Dunn’s multiple comparison test for non-parametric data. A p-value of less than 0.05 was considered statistically significant.

## Results

### Characterization of HDF spheroids

In this study, the Elplasia® microcavity plates were used for culturing 3D spheroids, offering a reproducible environment for generating size-controlled HDF spheroids. After a 4-day seeding period, spheroids were collected in culture flasks **(Figure 1A)**. The seeding cell density influences the size of spheroids, with higher cell numbers resulting in larger and more densely packed spheroids. HDF spheroids, each containing 125, 250, 500, 1,000, and 2,000 cells, were estimated to have an average diameter of 86.85±1.68, 95.3±1.81, 113.6±1.82, 124±2.2, and 151.6±2.4 μm **(Figure 1B)**, and an average cross-sectional area of 7435±247.4, 9570±307.8, 11545±376.7, 14367±352.9, and 20731±646.9 μm^2^, respectively **(Figure 1C)**. Following overnight incubation, individual fibroblasts migrated away from the spheroids and re-attached to the culture flasks **(Figure 1A, lower panels)**, suggesting fibroblasts inside the spheroids were viable and preserved their phenotypic features, exhibiting immunopositivity for markers commonly expressed in fibroblasts (e.g., αSMA and Vimentin) and MSCs (e.g., CD44 and CD90) **(Figure 1D)**.

**Fig. 1.**
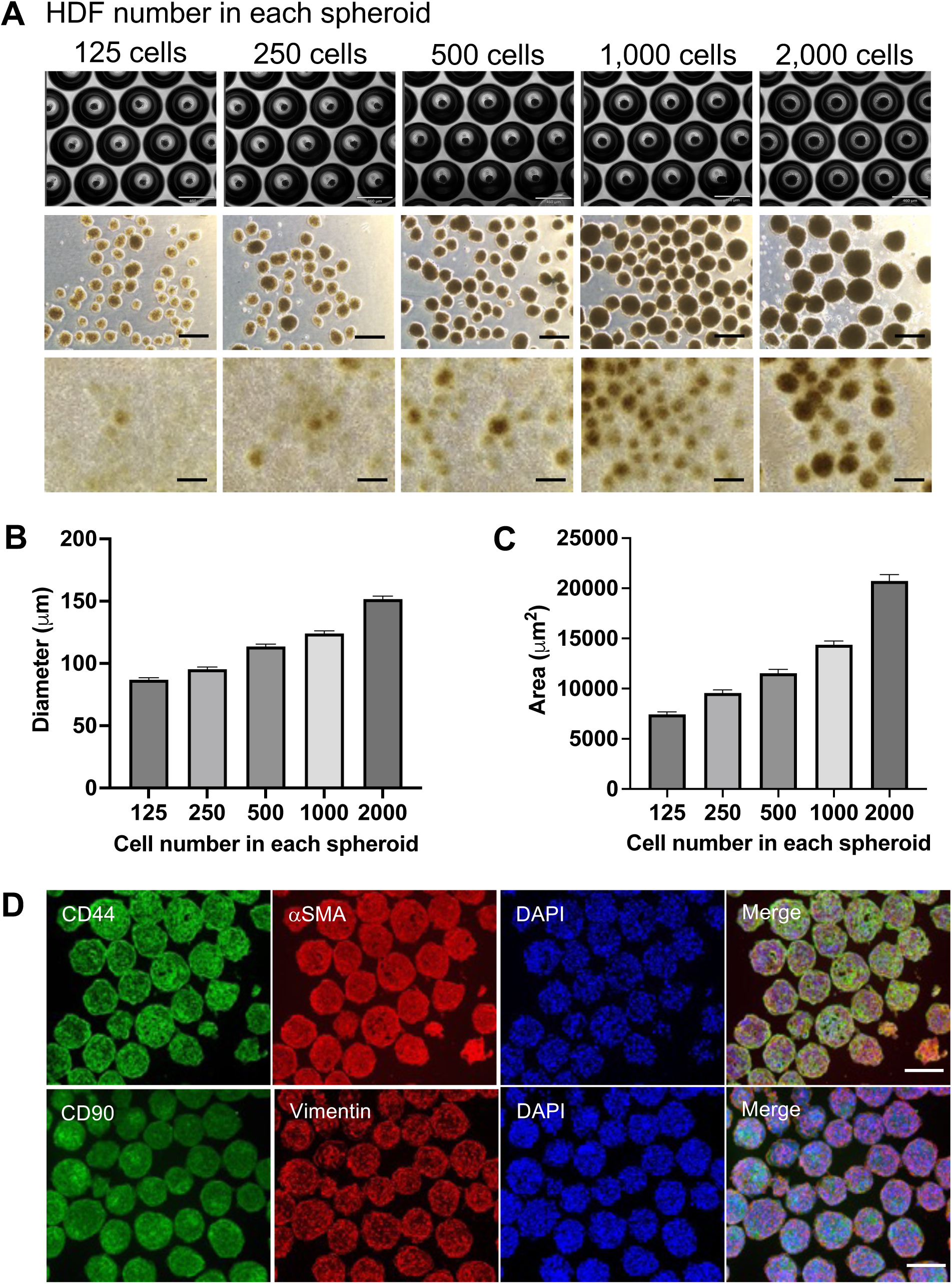
Morphological and phenotypic features of human dermal fibroblast (HDF) spheroids. (A) Row 1. Microscopic images of HDF spheroids containing various cell numbers in each spheroid in 24-well Elplasia® microcavity plates. **Row 2** After four days of culture, spheroids were harvested and imaged. **Row 3** Individual fibroblasts migrated away from the spheroids and re-attached to the culture plates upon overnight incubation. Scale bar: (Row 1) 460 μm; (Rows 2-3) 300 μm. Both (**B)** diameter (μm) and (**C)** cross-sectional area (μm^2^) of HDF spheroids increased with a higher cell number in each spheroid. Data shown are mean ± SEM. *n* = 45-54 per group. (**D)** Phenotypic features of HDF were maintained in spheroid cultures, as shown by immunofluorescent staining for CD44, α smooth muscle actin (αSMA), CD90, and Vimentin. Scale bar: 150 μm.

### Comparison of size, viability, and phenotypic features between HDF single cells and spheroids

Cells dissociated from HDF spheroids showed a high viability of over 97% after a 7-day incubation at 4°C, a significant improvement compared to the 68% viability in single cells from monolayer cultures **(Figure 2A)**. Although the diameter of spheroids appeared unaffected qualitatively during the 7-day incubation (data not shown), the individual cells dissociated from spheroids displayed a 30% decrease in diameter (from 18.73±0.23 μm to 13.13±0.18 μm, p<0.0001) **(Figure 2B)**. The smaller size of cells dissociated from spheroids could reflect significant changes in their structural and metabolic environments (21, 34). However, it is also possible that the reduced size is not intrinsic to the spheroid structure but rather an artifact caused by dissociation conditions, such as enzymatic treatment or mechanical disruption (35). Immunofluorescent staining showed an increase in the expression of S100A4 (p<0.001) and CD105 (p<0.05), along with a decrease in ITGA5 (p<0.05) in cells dissociated from spheroids **(Figure 2C-F)**.

**Fig. 2.**
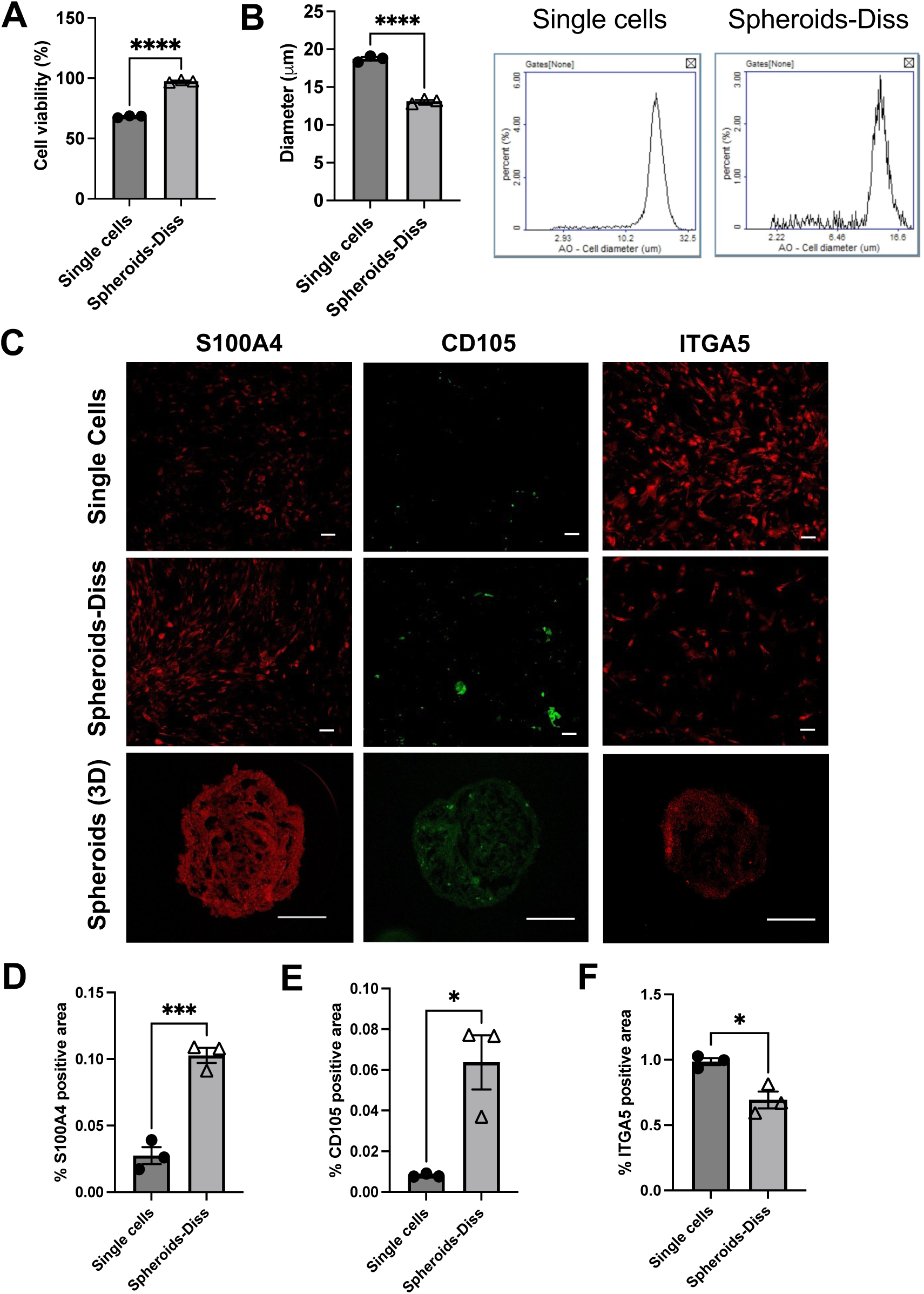
Comparison of size, phenotypic features, and viability between HDF single cells and spheroids. **(A)** Cells dissociated from HDF spheroids (Spheroids-Diss) maintained over 97% viability after a 7-day incubation at 4°C, compared to 68% in single cells. (**B)** Left: Cells dissociated from HDF spheroids exhibited reduced size compared with single cells. Right: Representative images showing cell diameter measurements (x-axis, μm) through AO staining using the NucleoCounter NC-3000 instrument. Single-cell diameter falls within the range of 2.93-32.5 μm, while Spheroid-Diss falls within the range of 2.22-16.6 μm. (**C-F)** Immunofluorescent staining showed an increase in the expression of S100A4 and CD105, along with a decrease in integrin α5 (ITGA5) in cells dissociated from HDF spheroids. The immunopositivity was normalized to the immunopositive area for DAPI. Scale bar: 100 μm. *n* = 3 per group. Data shown are mean ± SEM. *p<0.05, ***p < 0.001, and ****p < 0.0001.

### HDF spheroids alleviated the severity of IMQ-induced psoriasis

In this mouse model of psoriasis, daily IMQ application led to a gradual decline in body weight, resulting in an approximately 10% decrease on Day 4 in the IMQ control group. HDF spheroid treatment did not significantly alter this trend **(Figure 3B and Figure 4B)**. Clinical scores for thickness, erythema, and scaling exhibited a progressive increase in psoriatic skin lesions, reaching the highest average PASI score of 3.4 on Day 8 in the IMQ control group **(Figure 3 C-G)**. A single administration of HDF spheroids on Day 4 (PASI=2) significantly reduced skin thickness on Days 7 and 8 and reduced psoriasis severity from Day 6 to Day 8, with a remarkable 35% decrease in average PASI on Day 8 (from 3.4±0.11 to 2.2±0.13; p<0.0001) **(Figure 3 D-G)**. Topical application of IMQ to the back skin resulted in splenomegaly, evidenced by increased spleen weight and length (p<0.0001) compared to healthy control mice. HDF spheroids significantly attenuated the IMQ-induced increase in spleen weight and length **(Figure 3 H-J)**. IMQ-induced psoriasis led to a significant reduction in RBC (p<0.01), hemoglobin (p<0.01), and platelets (p<0.05) **(Figure 3 K-M)**, and alterations in white blood cells (WBC) differential count, particularly monocytes in the peripheral blood **(Figure 3 N, O)**. A single administration of HDF spheroids ameliorated these effects, restoring RBC, hemoglobin, platelet, and monocyte levels to those comparable to the healthy controls **(Figure 3 K-O)**.

**Fig. 3.**
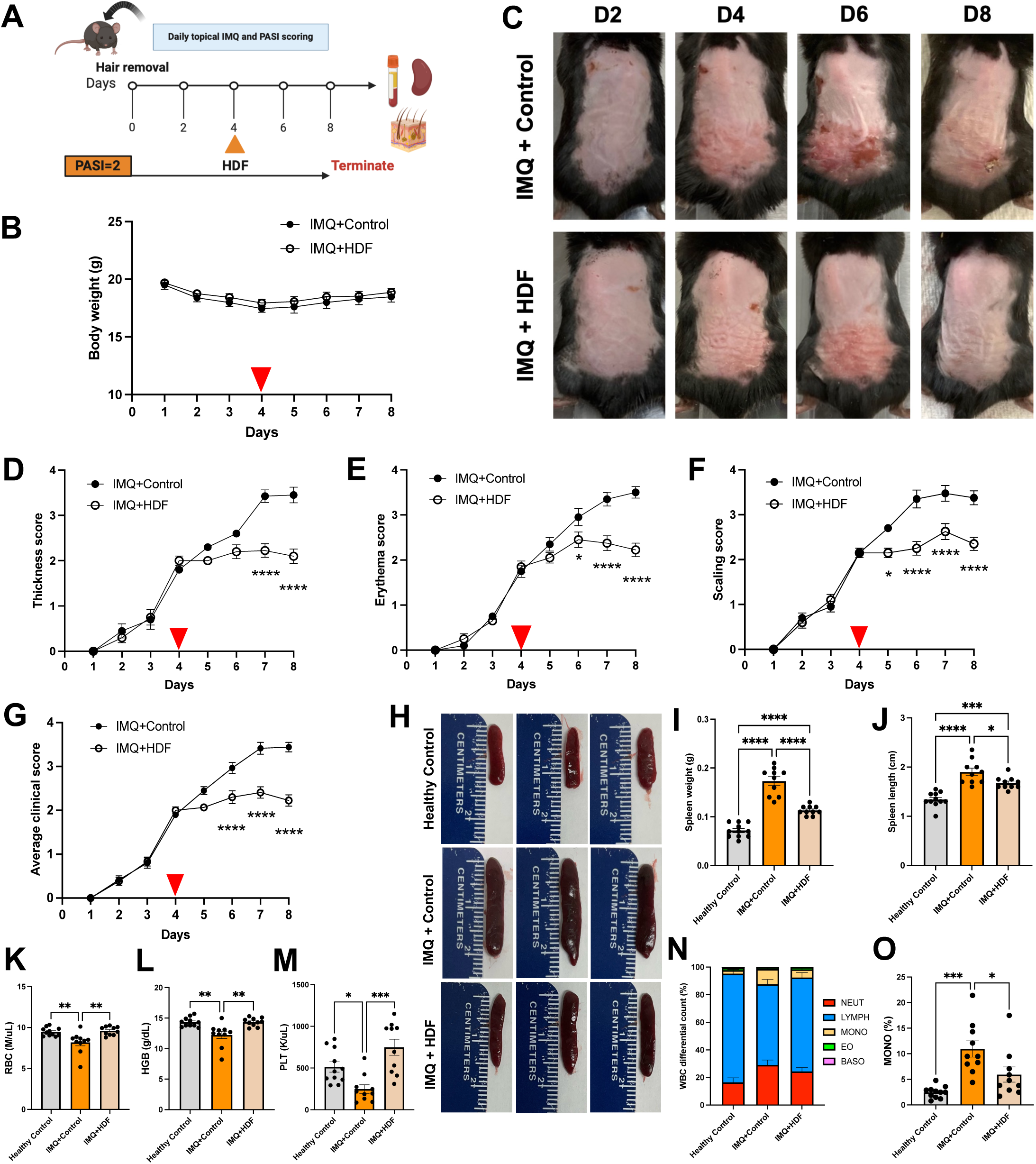
A single administration of HDF spheroids alleviated mild psoriasis. **(A)** The experimental timeline involved topically administering 31.25 mg of 5% IMQ to the lower dorsal skin of 10-week-old C57BL/6J female mice. Daily assessments of disease severity were conducted using the Psoriasis Area and Severity Index (PASI). A vehicle solution with (IMQ+HDF) or without HDF spheroids (IMQ+Control) was intravenously injected via the tail vein when the average PASI score reached 2 (Day 4), and euthanasia occurred when the IMQ+Control group reached a PASI score of 3-4 (Day 8). (**B)** Representative images of mouse back skin after a single administration of HDF spheroids. (**C)** Mouse body weight was monitored daily. The red triangle indicates the time of HDF spheroid intravenous injection. Daily monitoring included (**D-F)** PASI scores for thickness, erythema, and scaling on a scale of 0 (no incidence of disease) to 4 (most severe), respectively, and (**G)** the average PASI score for each animal. (**H)** Topical IMQ application induced splenomegaly, as indicated by increases in the spleen (**I)** weight and (**J)** length compared with healthy controls. HDF spheroids significantly reduced the IMQ-induced increase in spleen weight and length. HDF spheroids ameliorated the effects of IMQ, restoring **(K)** red blood cell (RBC), **(L)** hemoglobin (HGB), **(M)** platelet (PLT) counts, **(N)** % of WBC differential count (NEUT, neutrophils; LYMPH, lymphocytes; MONO, monocytes; EO, eosinophils; BASO, basophils), and **(O)** monocyte levels to those comparable to the healthy controls. Data shown are mean ± SEM. *n* =10-11 per group. *p<0.05, **p<0.01, ***p < 0.001, and ****p < 0.0001.

**Fig. 4.**
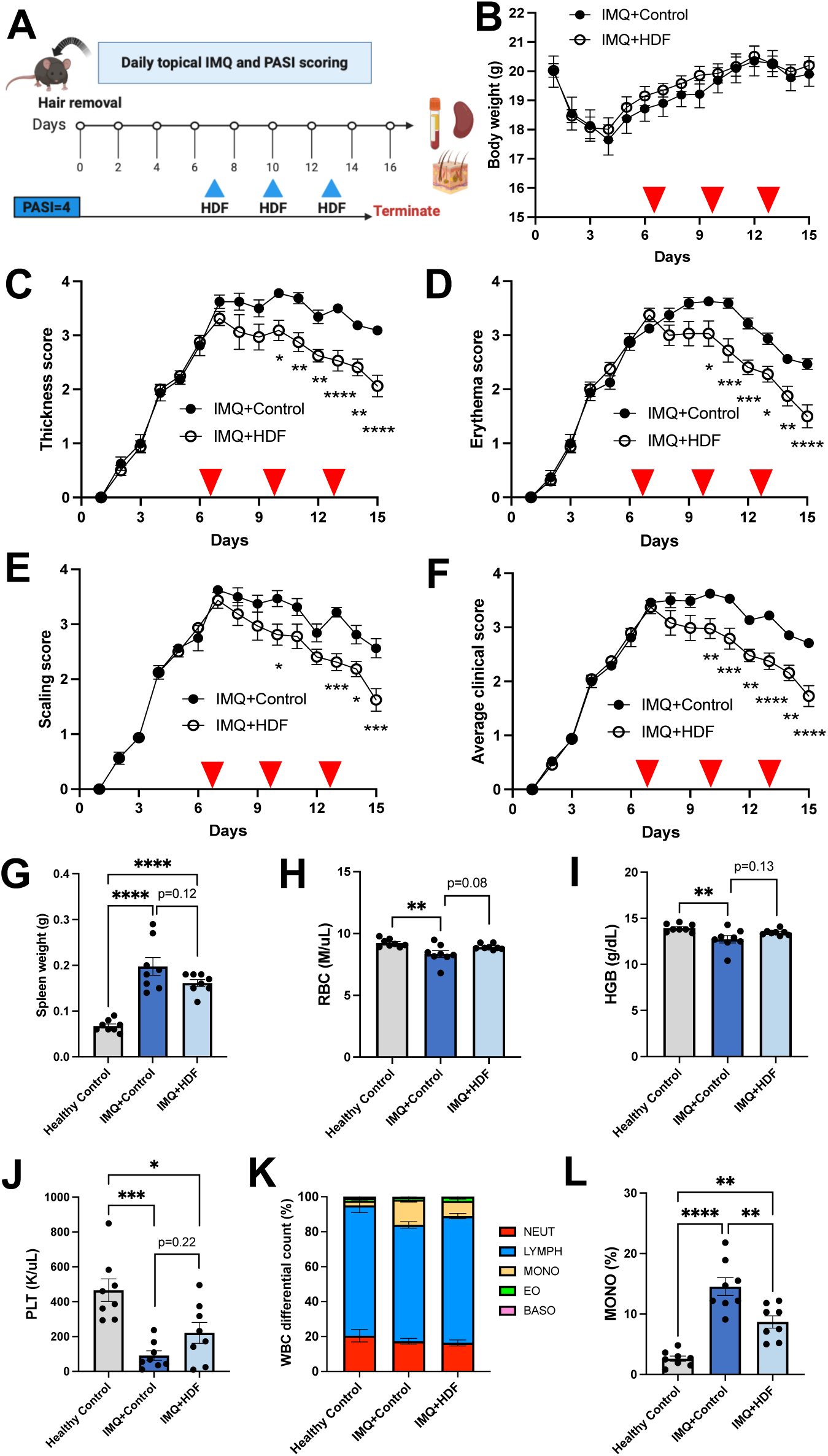
Multiple administrations of HDF spheroids alleviated moderate-to-severe psoriasis. **(A)** Psoriasis was induced and maintained by daily topical application of IMQ. HDF spheroids were intravenously injected when the average PASI score reached 3-4 (Day 7) and repeated on Days 10 and 13, and euthanasia occurred on Day 15. (**B)** Mouse body weight. The red triangle indicates the time of HDF spheroid intravenous injection. Daily monitoring included (**C-E)** PASI scores for thickness, erythema, and scaling and (**F)** the average PASI score for each animal. (**G)** IMQ-treated groups exhibited IMQ-induced splenomegaly. Multiple administrations of HDF spheroids induced a trend towards reduced spleen mass in psoriatic mice, but not significantly. An improvement in **(H)** RBC, (**I)** HGB, and (**J)** PLT was observed in psoriatic mice treated with multiple HDF spheroids, but not significantly. (**K, L)** Analysis of the differential count of WBC revealed trends consistent with the therapeutic effects of a single administration of HDF spheroids in mild psoriasis. A significant decrease in monocytes in the peripheral blood was observed in psoriatic mice treated with HDF spheroids. Data shown are mean ± SEM. *n* =8-11 per group. *p<0.05, **p<0.01, ***p < 0.001, and ****p < 0.0001.

In the multiple administration study, psoriatic lesions were sustained through daily IMQ application. As expected, the PASI scores for thickness, erythema, and scaling peaked on Days 7-8, regardless of some inter-individual variability **(Figure 4 C-F)**. The initial injection was administered on Day 7, followed by repeated doses every 3 days on Days 10 and 13. In mice with moderate-to-severe psoriasis, multiple administrations of HDF spheroids led to a notable reduction in the average PASI by 18% on Day 10 (p<0.01), 26% on Day 13 (p<0.0001), and 36% on Day 15 (p<0.0001) compared to IMQ-only controls **(Figure 4F)**. While both IMQ-treated groups exhibited IMQ-induced splenomegaly, multiple administrations of HDF spheroids showed modest effects on reducing spleen weight, although not significantly (p=0.12) **(Figure 4G)**. HDF spheroids did not significantly improve RBC (p=0.08), hemoglobin (p=0.13), and platelets (p=0.22) in psoriatic mice when compared to IMQ-only controls **(Figure 4 H-J)**. This partial restoration could be attributed to the prolonged impact of IMQ on the immune system. However, HDF spheroids exhibited significant inhibitory effects on WBC differential count, particularly monocytes (p<0.01 compared to IMQ-only controls) **(Figure 4 K, L)**, which was also observed in mice with mild psoriasis with a single administration of HDF spheroids.

### HDF spheroids reduced epidermal thickening, immune cell infiltration, and cell proliferation in psoriatic lesions

Psoriatic skin lesions induced by IMQ display notable characteristics such as markedly thickened epidermis, acanthosis, and hyperkeratosis. Histological examination via H&E staining revealed a substantial 4.8-fold increase in epidermal thickness within the lesions (p<0.0001) compared to healthy controls. A single administration of HDF spheroids significantly mitigated this effect, resulting in a remarkable reduction in thickness, compared to IMQ-only controls from 131.94 ± 3.52 μm to 74 ± 4.32 μm (p<0.0001) **(Figure 5 A, B)**. Additionally, HDF spheroid treatment elicited a notable downregulation of immune cell markers associated with T lymphocytes (CD3) and macrophages (F4/80) in the dermis, as well as the cell proliferation marker (Ki67) in the epidermis within the lesions **(Figure 5 A, C-E)**.

**Fig. 5.**
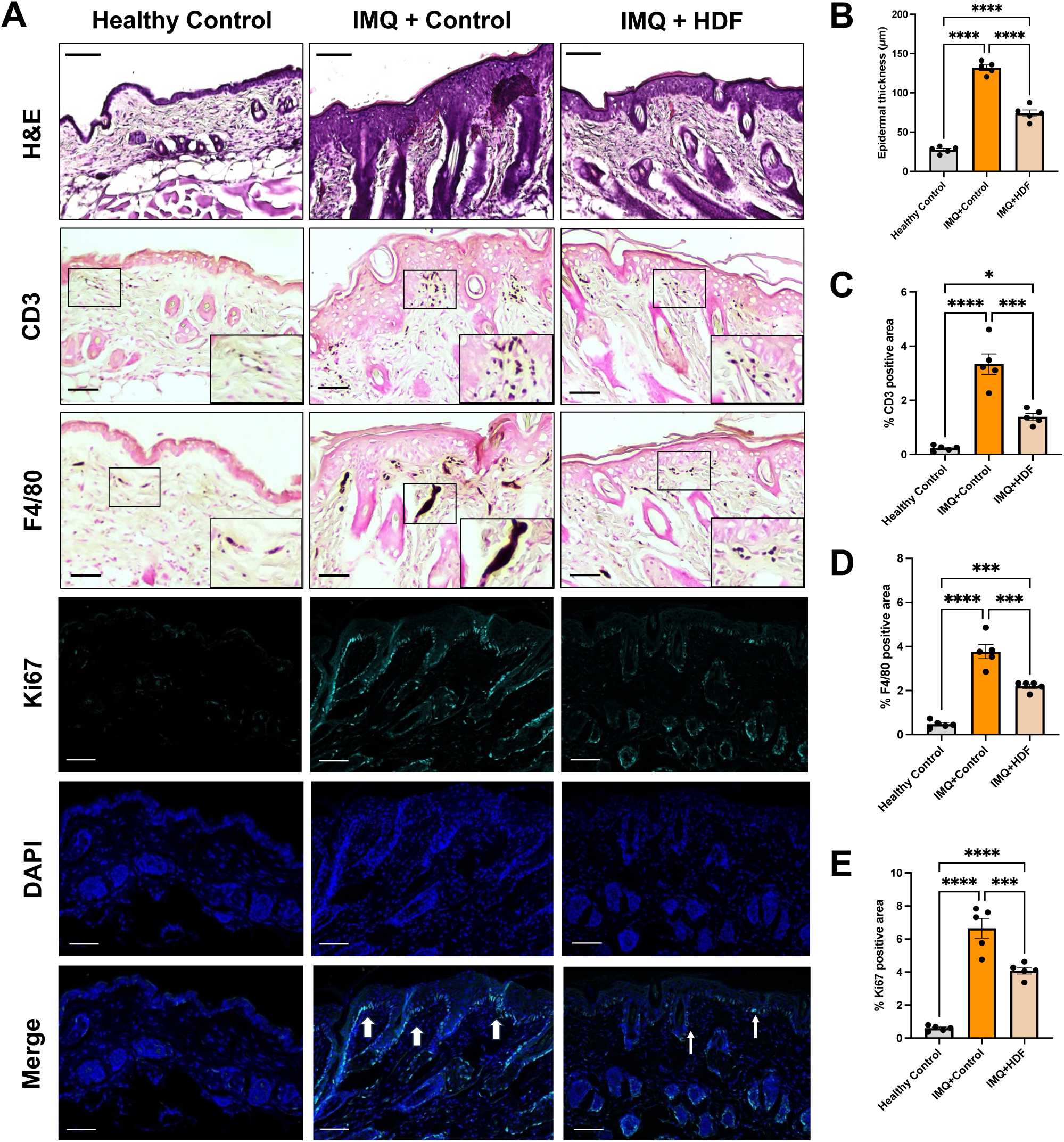
HDF spheroids reduced epidermal thickening, immune cell infiltration, and cell proliferation in psoriatic lesions. Representative images of **(A)** skin H&E and immunohistological staining for CD3 (T lymphocytes), F4/80 (macrophages), and Ki67 (cell proliferation) in Healthy Control, IMQ+Control, and IMQ+HDF mice. IMQ treatment induced psoriatic lesions with markedly thickened epidermis, acanthosis, hyperkeratosis, immune cell infiltration into the dermis, and elevated proliferation of keratocytes in the epidermis. **(B-E)** Treatment with HDF spheroids significantly inhibited epidermal thickening and resulted in a down-regulation of CD3, F4/80, and Ki67 in psoriatic lesions, as measured by % of immunopositive area using ImageJ. Scale bar: H&E 100 μm; CD3, F4/80, Ki67, DAPI, and merge 50 μm. Data shown are mean ± SEM. *n* = 5 per group. *p<0.05, ***p < 0.001, and ****p < 0.0001.

### Comparative analysis of therapeutic efficacy between HDF spheroids and anti-IL-23 treatment in psoriasis

Both anti-IL-23 and HDF spheroid treatments effectively reduced IMQ-induced psoriasis over the 7-day monitoring period **(Figure 6 B-F)**. The anti-IL-23 group showed an immediate and sustained reduction in disease severity, whereas the HDF spheroids-treated group exhibited a more gradual response (Days 7 and 8) **(Figure 6 C-F)**. This difference in responses is likely related to both mode of action and frequency of treatments. Overall, both treatments provided effective relief from psoriasis, but their timelines of action differ.

**Fig. 6.**
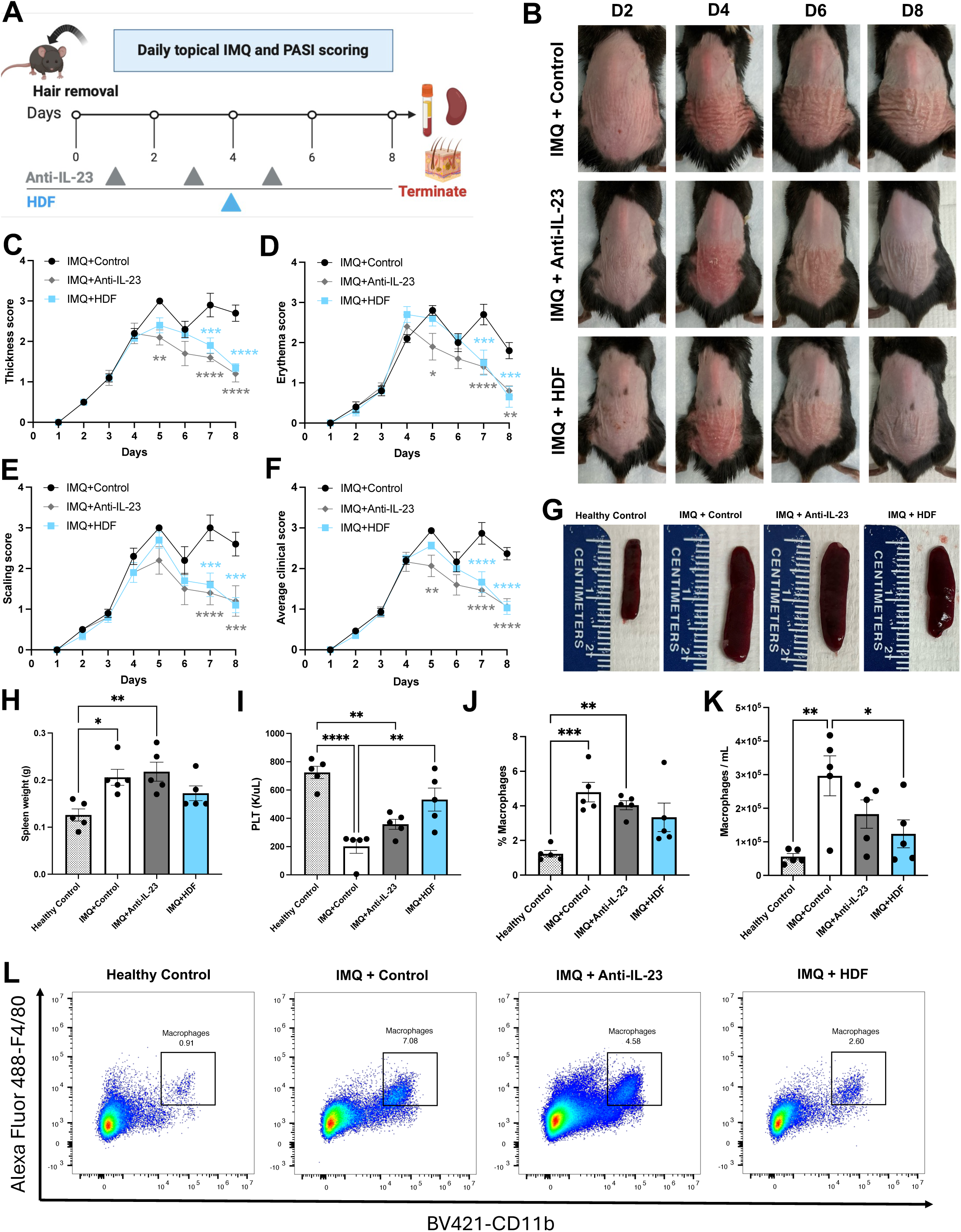
Comparative analysis of therapeutic efficacy between HDF spheroids and anti-IL-23 treatment in psoriasis. **(A)** Psoriasis was induced and maintained by daily topical application of IMQ. Anti-IL-23 was administered intraperitoneally on Days 1, 3, and 5 (IMQ+Anti-IL-23). HDF spheroids (IMQ+HDF) were intravenously injected when the average PASI score reached 2 (Day 4), and euthanasia occurred on Day 8. (**B)** Representative images of mouse back skin. Daily monitoring included (**C-E)** PASI scores for thickness, erythema, and scaling and (**F)** the average PASI score for each animal. (**G, H)** Topical IMQ-induced splenomegaly was still present after repeated administration of anti-IL-23 but was absent in the HDF spheroids-treated group. **(I)** An improvement in platelet counts was observed in psoriatic mice treated with HDF spheroids. **(J-L)** In flow cytometry analysis of isolated mouse splenocytes, cells were gated for macrophages using CD11b^+^ and F4/80^+^ markers. HDF spheroids significantly reduced macrophage infiltration in the spleen compared to IMQ-treated controls. Anti-IL-23 treatment did not exhibit a similar reduction in splenic macrophages. Data shown are mean ± SEM. *n* =5 per group. *p<0.05, **p<0.01, ***p < 0.001, and ****p < 0.0001.

Psoriasis mice treated with HDF spheroids showed no splenomegaly and exhibited improved platelet counts following the treatment. In contrast, mice treated with anti-IL-23 did not demonstrate these improvements, as splenomegaly was still present, and platelet counts were significantly lower than those of the healthy controls **(Figure 6 H, I)**. These findings suggest that HDF spheroids effectively alleviate spleen enlargement and enhance platelet recovery, unlike anti-IL-23 therapy. Flow cytometry analysis of isolated mouse splenocytes revealed a marked reduction in the population of macrophages (CD11b+F4/80+ cells) in HDF spheroids-treated mice. Other immune cells, such as DCs, B cells, T cells, and NK cells, did not change significantly (data not shown). This reduction, together with the absence of splenomegaly in these animals, could indicate that HDF spheroids might suppress macrophage infiltration in the spleen. In contrast, anti-IL-23-treated mice showed no significant reduction in macrophage populations and continued to exhibit splenomegaly **(Figure 6 J-L)**.

HDF spheroids and anti-IL-23 were both efficacious in inhibiting IMQ-induced epidermal thickening and reducing immune cell infiltration **(Figure 7A-E)**. Both treatment groups demonstrated fewer macrophages in skin lesions compared to healthy controls. However, the difference between the anti-IL-23-treated and healthy controls approached statistical significance (p=0.06, **Figure 7D**), suggesting that HDF spheroids were slightly more effective in reducing macrophage infiltration in the skin. Both HDF spheroids and anti-IL-23 treatments significantly inhibited systemic IL-17F **(Figure 7F)**, a key cytokine involved in psoriasis pathogenesis. Notably, HDF spheroids also increased IL-10 in the skin, restoring levels to those comparable to healthy controls. In contrast, anti-IL-23 treatment did not elevate IL-10 levels **(Figure 7G)**, highlighting the possible additional immunoregulatory benefit of HDF spheroids in promoting an anti-inflammatory local micro-environment.

**Fig. 7.**
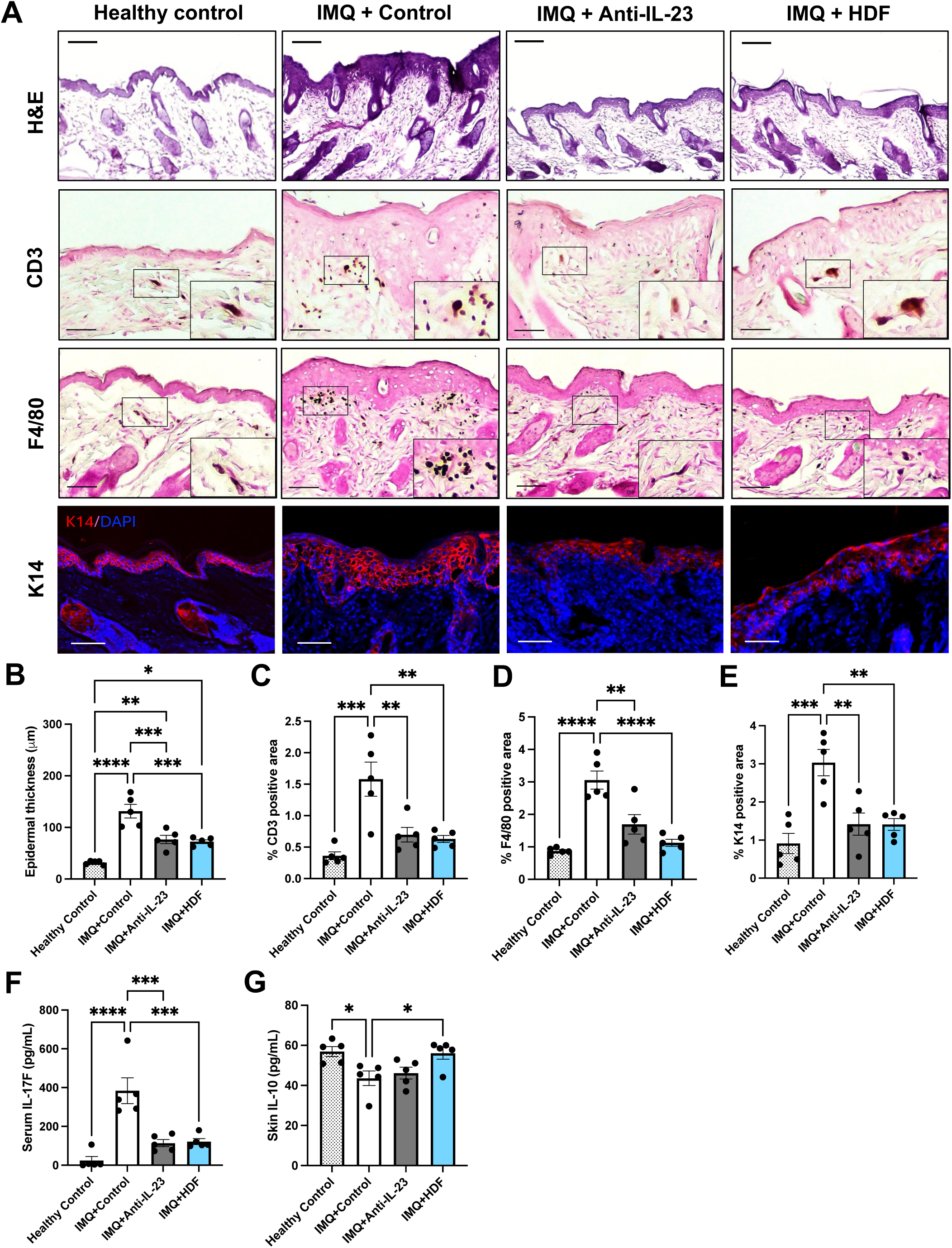
Histological assessment of skin lesions and cytokine profiling following HDF spheroids and anti-IL-23 treatment. Representative images of **(A)** skin H&E and immunohistological staining for CD3 (T lymphocytes), F4/80 (macrophages), and keratin 14 (K14, proliferative basal keratinocytes) in Healthy Control, IMQ+Control, IMQ+Anti-IL-23, and IMQ+HDF mice. **(B-E)** HDF spheroids and anti-IL-23 demonstrated comparable efficacy in inhibiting IMQ-induced epidermal thickening and reducing immune cell infiltration, as indicated by the down-regulation of CD3, F4/80, and K14. Scale bar: H&E 100 μm; CD3, F4/80, and K14 50 μm. **(F)** HDF spheroids and anti-IL-23 treatment significantly inhibited systemic IL-17F cytokine levels. **(G)** HDF spheroids restored IL-10 secretion in the skin to levels comparable to healthy controls, whereas anti-IL-23 did not. Data shown are mean ± SEM. *n* = 5 per group. *p<0.05, **p<0.01, ***p < 0.001, and ****p < 0.0001.

### Absence of IBMIR and toxicity in intravenous HDF spheroid administration

In this study, the risk of IBMIR was evaluated by administering HDFs intravenously in mice as either single cells or spheroids, followed by monitoring immune cell and coagulation-related changes at 1 hour, 24 hours, 3 days, and 7 days post-injection. The analysis revealed a time-dependent reduction in platelet levels and mean platelet volume (MPV) post HDF single-cell administration but not HDF spheroids **(Supplemental Figure 1 A, B)**. HDF single-cell administration rapidly influenced coagulatory indicators, with significant decreases in platelet counts observed within the first hour **(Supplemental Figure 1A)**. This reduction was followed by an increase in MPV **(Supplemental Figure 1B)**, likely due to platelet exhaustion and compensatory increased production by the bone marrow. Additionally, changes in immune cell counts were noted, with a strong trend towards decreased WBC and lymphocytes observed 24 hours post-administration **(Supplemental Figure 1 C, D)**. By Day 7, the levels of these coagulation indicators and immune cells gradually returned to baseline. In comparison, HDF spheroids did not induce significant IBMIR, as these parameters were comparable to those observed in vehicle-treated mice **(Supplemental Figure 1 A-D)**.

Following this, we assessed the acute (2-week) and chronic (12-week) safety/toxicity in mice with intravenously administered HDF spheroids (1×10^6^ cells/mouse, ∼150 µm). No mortality was observed over the 2-week and 12-week study periods. Analysis of relative tissue weights (heart, lungs, kidneys, liver, spleen, and brain) showed no significant differences between the control and HDF spheroid groups **(Supplemental Figure 1 E-J)**. CBC analysis and liver and kidney function biomarkers did not show clinically relevant changes between the two groups **(Supplemental Table 1)**. Histological examination of mouse lungs demonstrated no appreciable microscopic changes that could be attributed to HDF spheroid injection, such as thrombosis, fibrosis, or immune cell infiltration, with both control and HDF spheroids-treated mice exhibiting normal collagen deposition around bronchioles, indicating minimal adverse effects within the tested parameters **(Supplemental Figure 1K)**. These findings suggest that the administered dosage and size of HDF spheroids (1×10^6^ cells/mouse, ∼150 µm) are well tolerated over the evaluated timeframes.

## Discussion

This study demonstrated that HDF spheroids significantly alleviated psoriatic skin lesions, evidenced by reduced PASI scores, decreased epidermal thickening, and diminished immune cell infiltration. In addition to improving skin pathology, HDF spheroids also mitigated IMQ-induced splenomegaly and normalized blood cell parameters, such as RBC, hemoglobin, and platelet counts. Notably, the therapeutic efficacy of a single dose of HDF spheroids was comparable to that of multiple anti-IL-23 monoclonal antibody doses, a standard treatment for psoriasis. However, our study highlights the distinct advantage of HDF spheroids in enhancing immunomodulation, possibly through the inhibition of monocyte production and infiltration in the spleen and skin—an effect not observed with anti-IL-23. These findings suggest that HDF spheroids represent a promising alternative to traditional biological therapies for psoriasis.

Therapies using other cell-based approaches, such as MSCs, have demonstrated therapeutic effects in chronic inflammatory diseases, including psoriasis (36–38). However, the variability in MSC function, most often attributed to differences in tissue source, may lead to inconsistent therapeutic outcomes, presenting a major challenge for clinical translation (39, 40). In contrast, fibroblasts, which are more homogeneous in their biological functions and well-documented for their predictable immunosuppressive effects, offer a more consistent and reproducible therapeutic profile. Human dermal fibroblasts have demonstrated the ability to modulate lymphocyte activation, promote Tregs, and polarize macrophages toward an anti-inflammatory phenotype, making them a promising alternative to MSCs in conditions where consistent immune modulation is critical for long-term disease control (19). Fibroblasts have traditionally been recognized for their role in tissue repair, but recent evidence reveals their immunomodulatory properties. Fibroblasts can suppress immune activation through direct cell-cell interactions, the secretion of immunomodulatory molecules such as transforming growth factor-β (TGF-β), IL-6, prostaglandin E2 (PGE2) (41–43), and the downregulation of immune activation markers such as granzyme B and IL2RA (CD25) (44). Unlike antigen-presenting cells (APCs), fibroblasts have a limited capacity to stimulate lymphocytes compared to APCs, indicating a more regulatory function in immune responses than in directly activating T cells (44, 45). They also play a key role in shifting macrophages toward an anti-inflammatory phenotype by enhancing IL-10 and CD206 expression and reducing IL-12 (44), which is crucial for immune homeostasis in conditions like psoriasis, where immune dysregulation drives disease progression.

The use of 3D culture systems addresses key limitations of cell therapy by enhancing cell viability, reducing apoptosis, and improving the secretion of immunomodulatory factors (21, 35). In 2D cultures, cells injected intravenously often experience a significant loss due to the “first-pass effect,” where a sizable portion of cells become trapped in the lungs and are subsequently eliminated through phagocytosis (34). This results in a decline in viable cells and diminished therapeutic efficacy. In contrast, 3D culture systems better mimic *in vivo* conditions, allowing cells to maintain more natural interactions and functionalities while providing a protective microenvironment that enhances cell survival (46). For instance, a fluorescein-based cell labeling study has shown that spheroids persist in the lungs for up to 7 days post-injection, compared to 2D monolayer cells that nearly disappeared within 48 hours (23). The 3D culture also promotes cell survival by upregulating genes associated with resistance to hypoxia-induced apoptosis and reducing apoptosis overall (47). This improved survival supports the sustained release of immunomodulatory factors and contributes to a prolonged therapeutic effect. The close cell-to-cell contact within spheroids induces cell morphology and proliferation changes, including a reduction in cell size (35) that may facilitate the subsequent passage through the lungs and enhance therapeutic delivery to target tissues (34). Bartosh et al. demonstrated that cells from MSC spheroids maintained over 90% viability and were smaller than those from monolayer cultures, leading to fewer cells retained in the lungs and higher recovery rates in other tissues such as the liver, spleen, kidney, and heart (21).

In the present study, HDF spheroids similarly exhibited enhanced cell survival and smaller cell size compared to 2D cultures **(Figure 2 A, B)**. These characteristics may facilitate improved distribution of fibroblasts post-injection and lead to better therapeutic outcomes. The 3D culture method also promotes the upregulation of specific fibroblast and MSC markers **(Figure 2 C-E)**, suggesting that the spheroid environment enhances cell phenotypes and functionality. Consistent with our hypothesis, transcriptome analyses indicate that spheroid formation correlates with the up-regulation of genes associated with cytoskeletal dynamics, cell cycle regulation, and proliferation (21). Additionally, spheroid culture is associated with the down-regulation of ITGA5, a gene involved in pulmonary endothelial binding **(Figure 2 C, F)**. ITGA5 encodes a receptor for fibronectin and fibrinogen on pulmonary endothelial cells, and its decreased expression may reduce the fibroblasts’ ability to adhere to these ligands, potentially improving their migration from the lungs to target tissues (34). Our findings align with previous studies showing that ITGA5 expression is more prominent in 2D cultured cells, where integrin-mediated adhesion to flat surfaces and ECM proteins is needed. In contrast, cells cultured in spheroids rely more on cell-cell interactions and have altered adhesion dynamics compared to 2D cultures (34). In summary, these adaptive changes in fibroblast phenotypes may promote cell survival, reduce lung entrapment, and improve therapeutic efficacy. Further research is needed to elucidate the mechanisms underlying these effects and their functional implications for psoriasis treatment.

A critical challenge in cell therapy is the risk of IBMIR, which can substantially diminish the therapeutic efficacy of intravenously administered cells. IBMIR is characterized by the rapid activation of both coagulation and complement systems, triggering an inflammatory response that compromises cell survival and accelerates cell clearance (28, 48). Fibroblasts, especially when cultured as spheroids, may mitigate IBMIR through several mechanisms (21, 49, 50). The 3D structure of spheroids enhances the secretion of key anti-inflammatory mediators, such as TNF-stimulated gene/protein 6 (TSG-6), stanniocalcin-1 (STC1), and leukemic inhibitory factor (LIF) (21). This enhanced anti-inflammatory profile is likely linked to a reduction in the expression of pro-coagulant factors, thereby potentially lowering the risk of thrombosis. Moreover, the 3D configuration of spheroids may act as a physical barrier, shielding individual cells from direct exposure to blood components (50), which can limit the activation of coagulation and complement pathways associated with IBMIR. Our findings show that HDF spheroids do not significantly alter coagulation markers such as platelet counts and MPV following injection **(Supplemental Figure 1 A-D)**, aligning with previous studies demonstrating reduced thrombogenicity in 3D-cultured cells. By decreasing IBMIR and limiting coagulation activation, HDF spheroids may extend their immunomodulatory effects, ultimately improving therapeutic outcomes in inflammatory diseases like psoriasis. Further investigation into the mechanisms underlying this reduced IBMIR could enhance the safety and efficacy of fibroblast-based therapies.

Prior to assessing the therapeutic effects of HDF spheroids in a psoriasis mouse model, we conducted a pre-clinical safety evaluation of intravenous HDF spheroid injection. HDF spheroids were safe at the administered dose and size (1×10^6^ cells/mouse, 150 μm in average diameter), with no mortality or significant adverse effects observed in major tissues and organs over 2 and 12 weeks. Relative tissue weight (heart, lungs, kidneys, liver, spleen, and brain) was comparable between the control and HDF spheroids groups **(Supplemental Figure 1 E-J)**. Despite a few statistically significant but clinically negligible markers, CBC analysis, liver and kidney functions, and lung histology revealed no differences between the control and HDF spheroids groups **(Supplemental Table 1, Supplemental Figure 1K)**. The data obtained from the pre-clinical safety study can inform size and dosage optimization for fibroblast-based therapy, administration routes, and potential adverse effects, providing a critical safety profile that can be translated into clinical practice. By identifying and mitigating these risks in animal models, we can design safer and more effective clinical trials, increasing the likelihood of therapeutic outcomes in psoriatic patients. This approach ensures that potential complications are addressed early, ultimately improving patient safety and the therapeutic efficacy of fibroblast-based treatments in human clinical trials.

In the IMQ-induced psoriasis model, a single administration of HDF spheroids was sufficient to reduce disease severity in mice with mild psoriasis **(Figure 3)**, suggesting that early intervention may require fewer doses to restore immune balance. However, moderate-to-severe psoriasis is characterized by a higher degree of immune dysregulation, with elevated levels of pro-inflammatory cytokines (such as IL-17, IL-23, and TNF-α) and greater immune cell infiltration (51, 52). Multiple doses were necessary for moderate-to-severe psoriasis, highlighting the progressive nature of the disease and the need for repeated treatment to achieve sustained clinical improvement. In these cases, each additional dose of HDF spheroids led to a cumulative reduction in disease severity, as reflected by PASI score decreases of 18%, 26%, and 36% on Days 10, 13, and 15, respectively **(Figure 4)**. This cumulative benefit supports the idea that HDF spheroids exert their effects gradually, offering continued suppression of pro-inflammatory pathways over time. One limitation of this study was the regrowth of hair in C57BL/6J mice, which formed a physical barrier on the skin and reduced the effectiveness of IMQ application. This introduced variability in disease severity, especially in long-term studies assessing the therapeutic outcomes of HDF spheroids. In order to overcome this issue, future studies should consider using hairless mouse models, such as the SKH-1 strain (53), which allows for continuous IMQ application and a more reliable assessment of therapeutic efficacy.

Anti-IL-23 is an FDA-approved biologic therapy that specifically targets and inhibits IL-23, a cytokine involved in the inflammatory cascade of psoriasis. By neutralizing IL-23, this therapy effectively reduces Th17 cell activation and subsequent production of pro-inflammatory cytokines, thus alleviating local inflammation and psoriatic skin lesions (31). While both HDF spheroids and anti-IL-23 reduced psoriatic lesions and systemic IL-17F levels, HDF spheroids uniquely promoted IL-10 production within psoriatic skin lesions **(Figures 6 and 7)**, an anti-inflammatory cytokine associated with tissue repair (54). Histological analysis demonstrated that both treatments reduced epidermal thickness and immune cell infiltration, including CD3+ T cells and F4/80+ macrophages. Notably, the reduction in macrophage infiltration was more pronounced with HDF spheroids compared to anti-IL-23 therapy **(Figure 7D)**. Flow cytometry analysis of splenocytes revealed that HDF spheroids significantly decreased the macrophage population in the spleen, an effect not observed with anti-IL-23 treatment **(Figure 6 J-L)**. Therefore, we proposed that HDF spheroids mitigate psoriasis by modulating macrophage dynamics through two primary mechanisms: (1) enhancing the production of IL-10 in psoriatic skin, which promotes an anti-inflammatory environment, and (2) reducing the systemic and local macrophage population, thereby decreasing immune cell infiltration and epidermal thickening. In contrast, anti-IL-23 therapy focuses primarily on inhibiting Th17-mediated inflammation. This dual action on cytokine production and macrophage regulation suggests that HDF spheroids might shift the immune response from a pro-inflammatory state to a more regulated, anti-inflammatory state, ultimately leading to reduced psoriasis. Our findings highlight a broader immunomodulatory potential of HDF spheroids, which could improve outcomes for patients who do not respond adequately to current biologic therapies like anti-IL-23.

Despite these promising findings, certain challenges remain in translating HDF spheroids therapy into clinical investigations. One such challenge is the potential risk of microvascular embolism due to the size of the spheroids (55), though our safety studies have optimized spheroid size to mitigate this risk. In addition, understanding the long-term viability and migration patterns of the injected cells is critical for optimizing dosages and improving therapeutic outcomes in future clinical trials. Current methods for tracking cell fate, such as polymerase chain reaction (PCR) techniques for human-specific genes, immunofluorescent staining, and bioluminescence imaging, do not distinguish between viable and non-viable cells (34, 56, 57). A previous study isolating and re-culturing injected MSCs revealed a limited lifespan, with viable MSCs predominantly retained in the lungs for up to 24 hours post-injection (58). The lack of migration to other injured organs suggests that the detection of MSCs in other tissues may be from MSC debris or phagocytosed MSCs rather than viable ones, implying that the observed immunomodulatory effects of MSC are likely mediated by other cell types, ultimately influencing the immune response at target tissues. In this study, we demonstrate that HDF spheroids reduce psoriasis severity by modulating macrophage dynamics, which involves enhancing IL-10 production in psoriatic skin to create an anti-inflammatory environment and reduce both systemic and local macrophage populations. These findings suggest that HDF spheroids may offer a novel approach to managing psoriasis by influencing key inflammatory pathways. However, a global or tissue-specific monocyte-depletion model might be needed to fully understand and validate the effects to confirm HDF spheroids’ therapeutic potential and elucidate the precise mechanisms by which they modulate macrophage activity and contribute to disease amelioration.

### Conclusions

In conclusion, HDF spheroids represent a novel and promising approach to cell-based therapy for psoriasis. Their efficacy varies with disease severity, with single administrations proving effective for mild cases, while multiple doses are required for moderate-to-severe conditions. HDF spheroids demonstrate comparable efficacy to anti-IL-23, coupled with distinct immunomodulatory advantages. Notably, HDF spheroids are safe both short-term and long-term, potentially addressing challenges such as IBMIR associated with 2D-based cell therapy. The enhanced therapeutic benefits of HDF spheroids are likely attributable to their superior fibroblast/stem cell phenotypes, reduced pulmonary binding of cells dissociated from the spheroids due to their smaller size, and improved survival. These findings highlight the potential of HDF spheroids as a viable and effective alternative to existing treatments.

## Declarations

### Ethics approval and consent to participate

The animal study protocols used for this study were approved by the Institutional Animal Care and Use Committee at the K2Bio Lab (Houston, TX, USA) and StillMeadow, Inc. (Sugar Land, TX, USA).

### Consent for publication

Not applicable.

### Availability of data and materials

The datasets used and/or analysed during the current study are available from the corresponding author on reasonable request.

### Authors’ contributions

All authors participated in manuscript preparation and revision. H.K., C.F., and P.O.H. conceived and designed the experiments. C.F., I.B.E., and B.J. performed the experiment and collected the data. C.F. analyzed the data and wrote the manuscript. C.F., I.B.E., C.B., S.G., N.B., B.J., S.R., M.T., H.K., and P.O.H. discussed the results and commented on the manuscript.

## List of abbreviations

ABC: avidin-biotin-peroxidase
APC: antigen-presenting cells
BCA: bicinchoninic acid
CBC: complete blood count
DAB: 3,3′-diaminobenzidine
DC: dendritic cells
ECM: extracellular matrix
FBS: fetal bovine serum
HDF: human dermal fibroblasts
HGB: hemoglobin
IBMIR: instant blood-mediated inflammatory reaction
IL: interleukin
IMQ: imiquimod
ITGA5: integrin α5
LIF: leukemic inhibitory factor
MSC: mesenchymal stem cells
MPV: mean platelet volume
PASI: Psoriasis Area and Severity Index
PBS: phosphate-buffered saline
PCR: polymerase chain reaction
PLA: plasma-Lyte A
PLT: platelets
RBC: red blood cells
RT: room temperature
SEM: standard error of the mean
STC1: stanniocalcin-1
TNF-α: tumor necrosis factor α
Tregs: regulatory T cells
TSG-6: TNF-stimulated protein 6
WBC: white blood cells

## Acknowledgments

Not applicable.

## Figure legends

**Supplemental Fig. 1.** Pre-clinical safety evaluation of intravenous HDF spheroid injection in 10-week-old C57BL/6J female mice. Mice (n=5 per group) were intravenously administered with HDF cells as single cells (HDF single cells) or spheroids (HDF spheroids). The effects on coagulation-related factors and immune response were assessed at 1 hour, 24 hours, 3 days, and 7 days post-administration, including **(A)** platelets (K/uL), **(B)** mean platelet volume (MPV, K/uL), **(C)** WBC (K/uL), and **(D)** lymphocyte counts. HDF single-cell administration rapidly influenced coagulation factors, with significant decreases in platelet counts observed within the first hour and a subsequent increase in MPV. Changes in immune cell counts were noted, with a trend towards decreased WBC and lymphocytes 24 hours post-administration. By Day 7, the levels of these coagulation factors and immune cells gradually returned to baseline. When assessing the safety of intravenously administering HDF spheroids, mice (*n* = 4-5 per group) body weight, length, and relative tissue weight (%) of the heart, lungs, kidneys, liver, spleen, and brain in the **(E-G)** 2-week and **(H-J)** 12-week studies did not show statistical differences between the vehicle (Control) and HDF spheroid (HDF) groups. **(K)** Representative lung H&E and Masson’s trichrome (MT) images for acute (2-week) and chronic (12-week) toxicity. No microscopic changes were induced by intravenous injection of HDF spheroids in the mouse lungs. Control and HDF-treated mice demonstrated some collagen deposition (blue) around bronchioles (normal finding). Scale bar: 100 μm. Data shown are mean ± SEM. *p<0.05 and **p<0.01.

## References

1. Armstrong AW, Mehta MD, Schupp CW, Gondo GC, Bell SJ, Griffiths CEM. Psoriasis Prevalence in Adults in the United States. JAMA Dermatol. 2021;157(8):940–6.

2. Puig L. Cardiometabolic Comorbidities in Psoriasis and Psoriatic Arthritis. Int J Mol Sci. 2017;19(1).

3. Lee HJ, Kim M. Challenges and Future Trends in the Treatment of Psoriasis. Int J Mol Sci. 2023;24(17).

4. Parisi R, Iskandar IYK, Kontopantelis E, Augustin M, Griffiths CEM, Ashcroft DM. National, regional, and worldwide epidemiology of psoriasis: systematic analysis and modelling study. Bmj. 2020;369:m1590.

5. Rendon A, Schäkel K. Psoriasis Pathogenesis and Treatment. Int J Mol Sci. 2019;20(6).

6. Jeon C, Sekhon S, Yan D, Afifi L, Nakamura M, Bhutani T. Monoclonal antibodies inhibiting IL-12, –23, and –17 for the treatment of psoriasis. Hum Vaccin Immunother. 2017;13(10):2247–59.

7. Raimondo A, Balato A, Megna M, Balato N. Limitations of current monoclonal antibodies for plaque-type psoriasis and an outlook for the future. Expert Opin Biol Ther. 2018;18(6):605–7.

8. Chen M, Peng J, Xie Q, Xiao N, Su X, Mei H, et al. Mesenchymal Stem Cells Alleviate Moderate-to-Severe Psoriasis by Reducing the Production of Type I Interferon (IFN-I) by Plasmacytoid Dendritic Cells (pDCs). Stem Cells Int. 2019;2019:6961052.

9. Lee YS, Sah SK, Lee JH, Seo KW, Kang KS, Kim TY. Human umbilical cord blood-derived mesenchymal stem cells ameliorate psoriasis-like skin inflammation in mice. Biochem Biophys Rep. 2017;9:281–8.

10. Sah SK, Park KH, Yun CO, Kang KS, Kim TY. Effects of Human Mesenchymal Stem Cells Transduced with Superoxide Dismutase on Imiquimod-Induced Psoriasis-Like Skin Inflammation in Mice. Antioxid Redox Signal. 2016;24(5):233–48.

11. Campanati A, Orciani M, Sorgentoni G, Consales V, Mattioli Belmonte M, Di Primio R, et al. Indirect co-cultures of healthy mesenchymal stem cells restore the physiological phenotypical profile of psoriatic mesenchymal stem cells. Clin Exp Immunol. 2018;193(2):234–40.

12. Zhao X, Jiao J, Li X, Hou R, Li J, Niu X, et al. Immunomodulatory effect of psoriasis-derived dermal mesenchymal stem cells on TH1/TH17 cells. Eur J Dermatol. 2021;31(3):318–25.

13. Ichim TE, O’Heeron P, Kesari S. Fibroblasts as a practical alternative to mesenchymal stem cells. J Transl Med. 2018;16(1):212.

14. Dermagraft: Use in the Treatment of Chronic Wounds. Advances in Wound Care. 2012;1(3):138–41.

15. Petrof G, Martinez-Queipo M, Mellerio JE, Kemp P, McGrath JA. Fibroblast cell therapy enhances initial healing in recessive dystrophic epidermolysis bullosa wounds: results of a randomized, vehicle-controlled trial. Br J Dermatol. 2013;169(5):1025–33.

16. Jalili RB, Zhang Y, Hosseini-Tabatabaei A, Kilani RT, Khosravi Maharlooei M, Li Y, et al. Fibroblast Cell-Based Therapy for Experimental Autoimmune Diabetes. PLoS One. 2016;11(1):e0146970.

17. Jalili RB, Kilani RT, Li Y, Khosravi-Maharlooie M, Nabai L, Wang EHC, et al. Fibroblast cell-based therapy prevents induction of alopecia areata in an experimental model. Cell Transplant. 2018;27(6):994–1004.

18. Bouffi C, Bony C, Jorgensen C, Noël D. Skin fibroblasts are potent suppressors of inflammation in experimental arthritis. Ann Rheum Dis. 2011;70(9):1671–6.

19. Ichim TE, O’Heeron P, Perez J, Liu P, Min W-P, Kesari S. Fibroblasts as an Alternative to Mesenchymal Stem Cells with Successful Treatment and Immune Modulation in EAE Model of Multiple Sclerosis. bioRxiv. 2020:2020.06.04.133249.

20. Wang X, Jiang B, Sun H, Zheng D, Zhang Z, Yan L, et al. Noninvasive application of mesenchymal stem cell spheres derived from hESC accelerates wound healing in a CXCL12-CXCR4 axis-dependent manner. Theranostics. 2019;9(21):6112–28.

21. Bartosh TJ, Ylöstalo JH, Mohammadipoor A, Bazhanov N, Coble K, Claypool K, et al. Aggregation of human mesenchymal stromal cells (MSCs) into 3D spheroids enhances their antiinflammatory properties. Proc Natl Acad Sci U S A. 2010;107(31):13724–9.

22. Jiang B, Yan L, Miao Z, Li E, Wong KH, Xu RH. Spheroidal formation preserves human stem cells for prolonged time under ambient conditions for facile storage and transportation. Biomaterials. 2017;133:275–86.

23. Shimazawa Y, Kusamori K, Tsujimura M, Shimomura A, Takasaki R, Takayama Y, et al. Intravenous injection of mesenchymal stem cell spheroids improves the pulmonary delivery and prolongs in vivo survival. Biotechnol J. 2022;17(1):e2100137.

24. Song YC, Park GT, Moon HJ, Choi EB, Lim MJ, Yoon JW, et al. Hybrid spheroids containing mesenchymal stem cells promote therapeutic angiogenesis by increasing engraftment of co-transplanted endothelial colony-forming cells in vivo. Stem Cell Res Ther. 2023;14(1):193.

25. Yan L, Jiang B, Niu Y, Wang H, Li E, Yan Y, et al. Intrathecal delivery of human ESC-derived mesenchymal stem cell spheres promotes recovery of a primate multiple sclerosis model. Cell Death Discov. 2018;4:28.

26. Jiang B, Fu X, Yan L, Li S, Zhao D, Wang X, et al. Transplantation of human ESC-derived mesenchymal stem cell spheroids ameliorates spontaneous osteoarthritis in rhesus macaques. Theranostics. 2019;9(22):6587–600.

27. van der Fits L, Mourits S, Voerman JS, Kant M, Boon L, Laman JD, et al. Imiquimod-induced psoriasis-like skin inflammation in mice is mediated via the IL-23/IL-17 axis. J Immunol. 2009;182(9):5836–45.

28. Moll G, Ankrum JA, Kamhieh-Milz J, Bieback K, Ringdén O, Volk HD, et al. Intravascular Mesenchymal Stromal/Stem Cell Therapy Product Diversification: Time for New Clinical Guidelines. Trends Mol Med. 2019;25(2):149–63.

29. Jabeen M, Boisgard AS, Danoy A, El Kholti N, Salvi JP, Boulieu R, et al. Advanced Characterization of Imiquimod-Induced Psoriasis-Like Mouse Model. Pharmaceutics. 2020;12(9).

30. Neu SD, Strzepa A, Martin D, Sorci-Thomas MG, Pritchard KA, Jr., Dittel BN. Myeloperoxidase Inhibition Ameliorates Plaque Psoriasis in Mice. Antioxidants (Basel). 2021;10(9).

31. Zhou L, Wang Y, Wan Q, Wu F, Barbon J, Dunstan R, et al. A non-clinical comparative study of IL-23 antibodies in psoriasis. MAbs. 2021;13(1):1964420.

32. Rizzo HL, Kagami S, Phillips KG, Kurtz SE, Jacques SL, Blauvelt A. IL-23-mediated psoriasis-like epidermal hyperplasia is dependent on IL-17A. J Immunol. 2011;186(3):1495–502.

33. Fujiyama S, Nakahashi-Oda C, Abe F, Wang Y, Sato K, Shibuya A. Identification and isolation of splenic tissue-resident macrophage sub-populations by flow cytometry. Int Immunol. 2019;31(1):51–6.

34. Yeung CK, Yan Y, Yan L, Duan Y, Li E, Huang B, et al. Preclinical safety evaluation and tracing of human mesenchymal stromal cell spheroids following intravenous injection into cynomolgus monkeys. Biomaterials. 2022;289:121759.

35. Grässer U, Bubel M, Sossong D, Oberringer M, Pohlemann T, Metzger W. Dissociation of mono– and co-culture spheroids into single cells for subsequent flow cytometric analysis. Ann Anat. 2018;216:1–8.

36. Jasim SA, Yumashev AV, Abdelbasset WK, Margiana R, Markov A, Suksatan W, et al. Shining the light on clinical application of mesenchymal stem cell therapy in autoimmune diseases. Stem Cell Research & Therapy. 2022;13(1):101.

37. Choi EW. Adult stem cell therapy for autoimmune disease. Int J Stem Cells. 2009;2(2):122–8.

38. Zhang B, Lai RC, Sim WK, Choo ABH, Lane EB, Lim SK. Topical Application of Mesenchymal Stem Cell Exosomes Alleviates the Imiquimod Induced Psoriasis-Like Inflammation. Int J Mol Sci. 2021;22(2).

39. Galipeau J, Sensébé L. Mesenchymal Stromal Cells: Clinical Challenges and Therapeutic Opportunities. Cell Stem Cell. 2018;22(6):824–33.

40. Zhou T, Yuan Z, Weng J, Pei D, Du X, He C, et al. Challenges and advances in clinical applications of mesenchymal stromal cells. J Hematol Oncol. 2021;14(1):24.

41. Teramoto K, Igarashi T, Kataoka Y, Ishida M, Hanaoka J, Sumimoto H, et al. Clinical significance of PD-L1-positive cancer-associated fibroblasts in pN0M0 non-small cell lung cancer. Lung Cancer. 2019;137:56–63.

42. Lee B, Lee SH, Shin K. Crosstalk between fibroblasts and T cells in immune networks. Front Immunol. 2022;13:1103823.

43. Musso A, Condon TP, West GA, De La Motte C, Strong SA, Levine AD, et al. Regulation of ICAM-1-mediated fibroblast-T cell reciprocal interaction: implications for modulation of gut inflammation. Gastroenterology. 1999;117(3):546–56.

44. Denu RA, Nemcek S, Bloom DD, Goodrich AD, Kim J, Mosher DF, et al. Fibroblasts and Mesenchymal Stromal/Stem Cells Are Phenotypically Indistinguishable. Acta Haematol. 2016;136(2):85–97.

45. Cappellesso-Fleury S, Puissant-Lubrano B, Apoil PA, Titeux M, Winterton P, Casteilla L, et al. Human fibroblasts share immunosuppressive properties with bone marrow mesenchymal stem cells. J Clin Immunol. 2010;30(4):607–19.

46. Urzì O, Gasparro R, Costanzo E, De Luca A, Giavaresi G, Fontana S, et al. Three-Dimensional Cell Cultures: The Bridge between In Vitro and In Vivo Models. Int J Mol Sci. 2023;24(15).

47. Ryu NE, Lee SH, Park H. Spheroid Culture System Methods and Applications for Mesenchymal Stem Cells. Cells. 2019;8(12).

48. Moll G, Ankrum JA, Olson SD, Nolta JA. Improved MSC Minimal Criteria to Maximize Patient Safety: A Call to Embrace Tissue Factor and Hemocompatibility Assessment of MSC Products. Stem Cells Transl Med. 2022;11(1):2–13.

49. Moll G, Alm JJ, Davies LC, von Bahr L, Heldring N, Stenbeck-Funke L, et al. Do cryopreserved mesenchymal stromal cells display impaired immunomodulatory and therapeutic properties? Stem Cells. 2014;32(9):2430–42.

50. Moll G, Rasmusson-Duprez I, von Bahr L, Connolly-Andersen AM, Elgue G, Funke L, et al. Are therapeutic human mesenchymal stromal cells compatible with human blood? Stem Cells. 2012;30(7):1565–74.

51. Nestle FO, Kaplan DH, Barker J. Psoriasis. N Engl J Med. 2009;361(5):496–509.

52. Lowes MA, Suárez-Fariñas M, Krueger JG. Immunology of psoriasis. Annu Rev Immunol. 2014;32:227–55.

53. Xu Q, Liu Z, Cao Z, Shi Y, Yang N, Cao G, et al. Topical astilbin ameliorates imiquimod-induced psoriasis-like skin lesions in SKH-1 mice via suppression dendritic cell-Th17 inflammation axis. J Cell Mol Med. 2022;26(4):1281–92.

54. Asadullah K, Sterry W, Stephanek K, Jasulaitis D, Leupold M, Audring H, et al. IL-10 is a key cytokine in psoriasis. Proof of principle by IL-10 therapy: a new therapeutic approach. J Clin Invest. 1998;101(4):783–94.

55. Baranovskii DS, Klabukov ID, Arguchinskaya NV, Yakimova AO, Kisel AA, Yatsenko EM, et al. Adverse events, side effects and complications in mesenchymal stromal cell-based therapies. Stem Cell Investig. 2022;9:7.

56. Preda MB, Neculachi CA, Fenyo IM, Vacaru AM, Publik MA, Simionescu M, et al. Short lifespan of syngeneic transplanted MSC is a consequence of in vivo apoptosis and immune cell recruitment in mice. Cell Death Dis. 2021;12(6):566.

57. Sanchez-Diaz M, Quiñones-Vico MI, Sanabria de la Torre R, Montero-Vílchez T, Sierra-Sánchez A, Molina-Leyva A, et al. Biodistribution of Mesenchymal Stromal Cells after Administration in Animal Models and Humans: A Systematic Review. J Clin Med. 2021;10(13).

58. Eggenhofer E, Benseler V, Kroemer A, Popp FC, Geissler EK, Schlitt HJ, et al. Mesenchymal stem cells are short-lived and do not migrate beyond the lungs after intravenous infusion. Front Immunol. 2012;3:297.

