## Supplementary figures and images for "Human Dermal Fibroblast-derived Spheroids Demonstrate Efficacious Immune Modulation in a Psoriasis Mouse Model"

### Supplemental Figure 1

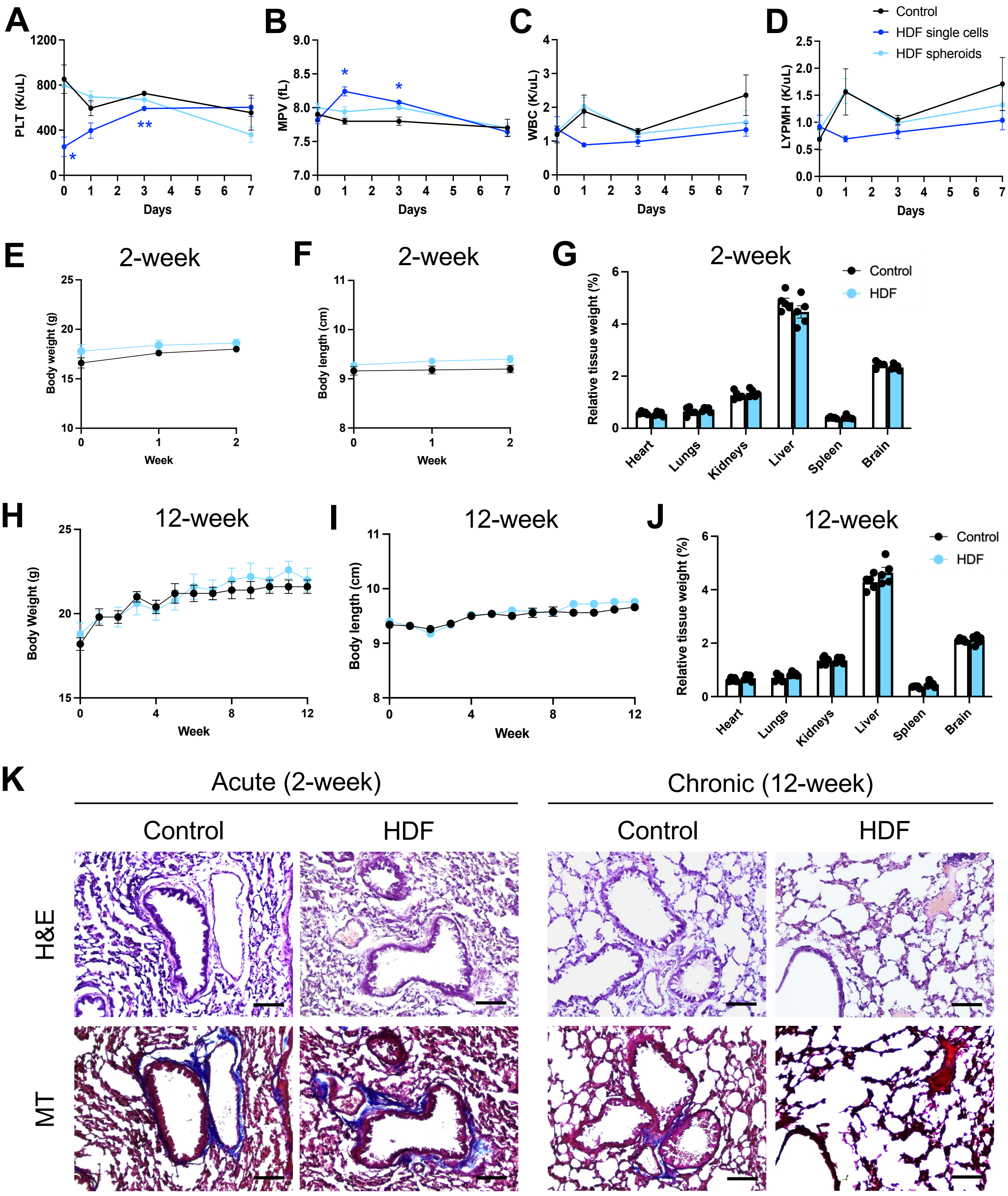
