## Supplemental Table 1 for "Human Dermal Fibroblast-derived Spheroids Demonstrate Efficacious Immune Modulation in a Psoriasis Mouse Model"

**Supplemental Table 1. Selected hematological and biochemical parameters of 10-week-old C57BL/6J female mice in the acute (2-week) and chronic (12-week) toxicity study.**

|  | **Acute toxicity  (2-week)** | | **Chronic toxicity  (12-week)** | |
| --- | --- | --- | --- | --- |
| **Parameters** | **Control** | **HDF** | **Control** | **HDF** |
| RBC  (x10^6^ cells /uL) | 9.43±0.32 | 9.14±0.22 | 10.02±0.93 | 10.3±0.54 |
| Hemoglobin (g/dL) | 14.42±0.56 | 14.18±0.39 | 15.66±1.37 | 15.05±0.79 |
| Hematocrit (%) | 51.36±2.31 | 49.06±1.39 | 50.6±4.9 | 48.53±3.42 |
| Platelet  (x10^3^ cells/uL) | 618.4±74.16 | 570.4±80.3 | 562.2±284.94 | 645.75±133.21 |
| WBC  (x10^3^ cells/uL) | 1.54±0.69 | 1.22±0.65 | 1.66±0.44 | 2.18±0.76 |
| Neutrophils  (x10^3^ cells/uL) | 0.15±0.05 | 0.15±0.1 | 0.43±0.11 | 0.55±0.27 |
| Lymphocytes  (x10^3^ cells/uL) | 1.34±0.62 | 1.01±0.59 | 1.16±0.41 | 1.52±0.75 |
| Monocytes  (x10^3^ cells/uL) | 0.03±0.02 | 0.03±0.02 | 0.05±0.02 | 0.05±0.01 |
| Eosinophils  (x10^3^ cells/uL) | 0.02±0.01 | 0.03±0.03 | 0.03±0.02 | 0.07±0.02* |
| Basophils  (x10^3^ cells/uL) | <0.01 | <0.01 | <0.01 | <0.01 |
| ALT (IU/L) | 24.79±2.59 | 22.91±3.99 | 32.13±4.78 | 25.93±2.65* |
| AST (IU/L) | 146.7±57.9 | 136.3±92.47 | 146.03±71.42 | 70.45±14.82 |
| CK (IU/L) | 629.33±257.65 | 516.59±449.93 | 466.79±318.06 | 146.26±63.38 |
| Creatinine (mg/dL) | 0.27±0.03 | 0.23±0.03 | 0.11±0.03 | 0.12±0.01 |
| LDH (IU/L) | 670.26±74.74 | 526.17±122.04 | 581.02±139.38 | 300.54±87.29** |
| TP (g/dL) | 4.62±0.13 | 4.37±0.25 | 4.49±0.36 | 4.368±0.28 |
| BUN (mg/dL) | 25.39±2.82 | 19.39±3.76* | 32.35±5.57 | 24.93±5.35 |
| Albumin (g/dL) | 3.18±0.12 | 2.98±0.13* | 3.01±0.26 | 3.09±0.11 |
| Globulin (g/dL) | 1.43±0.03 | 1.40±0.12 | 1.47±0.14 | 1.58±0.2 |
| A/G | 2.22±0.09 | 2.14±0.09 | 2.05±0.18 | 1.98±0.22 |

RBC, red blood cell; WBC, white blood cell; ALT, alanine transaminase; AST, aspartate transaminase; CK, creatine kinase; LDH, lactate dehydrogenase; TP, total protein; BUN, blood urea nitrogen; A/G, albumin/globulin ratio. Data shown are mean±SD. n=4-5 per group. *p<0.05 and **p<0.01.
